# Whole-brain connectivity atlas of glutamatergic and GABAergic neurons in mouse dorsal and median raphe nucleus

**DOI:** 10.1101/2020.12.15.422858

**Authors:** Zhengchao Xu, Zhao Feng, Mengting Zhao, Qingtao Sun, Lei Deng, Xueyan Jia, Tao Jiang, Pan Luo, Wu Chen, Jing Yuan, Xiangning Li, Hui Gong, Qingming Luo, Anan Li

## Abstract

The dorsal raphe nucleus (DR) and median raphe nucleus (MR) contain populations of glutamatergic and GABAergic neurons regulating diverse behavioral functions. Their whole-brain input-output circuits remain incompletely understood. We used viral tracing combined with fluorescence micro-optical sectioning tomography to generate a comprehensive whole-brain atlas of inputs and outputs of glutamatergic and GABAergic neurons in the DR and MR. We discovered that these neurons receive inputs from similar upstream brain regions. The glutamatergic and GABAergic neurons in the same raphe nucleus have divergent projection patterns with differences in critical brain regions. Specifically, MR glutamatergic neurons project to the lateral habenula via multiple pathways. Correlation and cluster analysis indicated that glutamatergic and GABAergic neurons in the same raphe nucleus receive inputs from heterogeneous neurons in upstream brain regions and send different collateral projections. This connectivity atlas provides insights into the cell heterogeneity, anatomical connectivity and behavioral functions of the raphe nucleus.

## Introduction

The dorsal raphe nucleus (DR) and median raphe nucleus (MR) are important modulatory centers involved in a multitude of functions (Domonkos et al.,2016; Huang et al., 2019; Szőnyi et al., 2019). They are implicated to have different and even antagonistic roles in the regulation of specific functions, such as emotional behavior, social behavior and aggression (Balázsfi et al., 2018; Ohmura et al., 2020; Teissier et al., 2015). The diverse regulatory processes are related to the connectivity of heterogeneous raphe neuron groups (Muzerelle et al., 2016; Nectow et al., 2017; Schneeberger et al., 2019). Deciphering precise neural circuits regarding the input and output circuits of cell-type-specific neurons in the raphe nucleus is fundamental for understanding their specific functions.

The DR and MR are heterogeneous and contain diverse types of neurons, including a large proportion of glutamatergic and GABAergic neurons (Huang et al., 2019; Pinto et al., 2019; Sos et al., 2017). Several studies have revealed that they are involved in specific functions. In the DR, glutamatergic neurons play an important role in reward processing (McDevitt et al., 2014; Liu et al., 2014), while GABAergic neurons are involved in regulating energy expenditure (Schneeberger et al., 2019), and they have opposite effects on feeding (Nectow et al., 2017). In the MR, glutamatergic neurons are critical for processing negative experiences, and activation of them could induce aversive behavior, aggression and depressive symptoms (Szőnyi et al., 2019). Furthermore, MR GABAergic neurons are involved in regulating hippocampal theta rhythm, which is crucial for learning and memory (Aitken et al., 2018; Li et al., 2005). The diverse functions of specific types of neurons in the raphe nucleus are highly dependent on their unique input-output circuits (Ren et al., 2018a). To have a more comprehensive understanding of the specific functions of glutamatergic and GABAergic neurons in the raphe nucleus, it is useful to get knowledge of how the cellular heterogeneity maps to whole-brain connectivity.

Previous studies have revealed that the DR and MR integrate massive inputs from and send outputs to many brain regions, such as the forebrain, hypothalamus and midbrain (Marcinkiewicz et al., 1989; Oh, et al., 2014; Peyron et al., 1997; Vertes et al., 2008). But these studies were unable to elucidate the neural connections of specific types of neurons. Studies using slice physiological recording combined with optogenetics found that DR GABAergic neurons receive long-range functional inputs from six upstream brain areas, including the prefrontal cortex, amygdala, lateral habenula (LH), lateral hypothalamic area (LHA), preoptic area and substantia nigra (Zhou et al., 2017). However, optogenetic technology and physiological recording usually focused on specific regions connected with targeted neurons, making it difficult to dissect whole-brain long-range connections. Genetic targeting of neuronal subpopulations with Cre driver mouse lines and virus tracing make it possible to label the whole-brain long-range connectivity of specific neurons (Callaway et al., 2015; Huang et al., 2013; Wickersham et al., 2007). Several studies have revealed a portion of the long-range connections of glutamatergic and GABAergic neurons in the DR and MR through viral tracing techniques. For instance, DR GABAergic neurons receive vast inputs and project to the dorsomedial nucleus of the hypothalamus (DMH) and bed nuclei of the stria terminalis (BST) (Schneeberger et al., 2019; Weissbourd et al., 2014). And MR glutamatergic neurons are innervated by certain aversion/fear or memory-related areas, such as the LH, and they innervate the LH, medial ventral tegmental area, medial septum and the vertical limbs of the diagonal bands of Broca (Szőnyi et al., 2019). Nevertheless, there is still a lack of whole-brain quantitative results and comprehensive analysis of the input-output circuits of glutamatergic and GABAergic neurons in the DR and MR. Furthermore, precise characterization and systematic quantitative analysis of whole-brain inputs and outputs require whole-brain high-resolution imaging of labeled neural structures and effective data processing methods to identify and integrate neural circuits.

In this study, we implemented a pipeline composed of viral tracing, whole-brain high-resolution imaging, data processing and analysis to dissect whole-brain inputs and outputs of glutamatergic and GABAergic neurons in the DR and MR and understand their organization principle. We used modified monosynaptic rabies viral tracers to label the input neurons and enhanced yellow fluorescent protein (EYFP)-expressing adeno-associated virus (AAV) to trace whole-brain axon projections. Combined with home-made fluorescence micro-optical sectioning tomography (fMOST) (Gong et al., 2016), we acquired whole-brain datasets of labeled inputs and outputs at single-neuron resolution. We identified the long-range input/output circuits, quantified the whole-brain distribution, analyzed the whole-brain connectivity pattern and generated a precise whole-brain atlas of inputs and outputs of glutamatergic and GABAergic neurons in the DR and MR, which could facilitate the understanding of their functional differences and provide anatomical foundations for investigating into their functions. Moreover, we developed the interactive website (http://atlas.brainsmatics.org/a/xu2011) to better present and share the raw data and results.

## Results

### Whole-brain mapping of monosynaptic input neurons to glutamatergic and GABAergic neurons in the DR and MR

To label the whole-brain inputs to glutamatergic and GABAergic neurons in the DR and MR, we used monosynaptic rabies tracing technique combined with Vglut2-Cre and Gad2-Cre driver line mice. First, Cre-dependent helper viruses, rAAV2/9-EF1α-DIO-His-TVA-BFP and rAAV2/9-EF1α-DIO-RG, were injected into the DR or MR. After three weeks, RV-ΔG-EnvA-GFP was injected into the same site (**Figure 1A**). The Cre-positive neurons infected by the Cre-dependent helper viruses could express the TVA receptor and glycoprotein. The rabies virus pseudotyped with the avian sarcoma leucosis virus glycoprotein EnvA could infect these neurons by binding TVA receptor specifically. Then, the rabies virus could be replenished with glycoprotein to retrogradely traverse to monosynaptic input neurons. The neurons co-labeled by GFP and BFP in the injection sites were starter cells, and GFP-labeled neurons were input neurons (**Figure 1B,C**).

**Figure 1.**
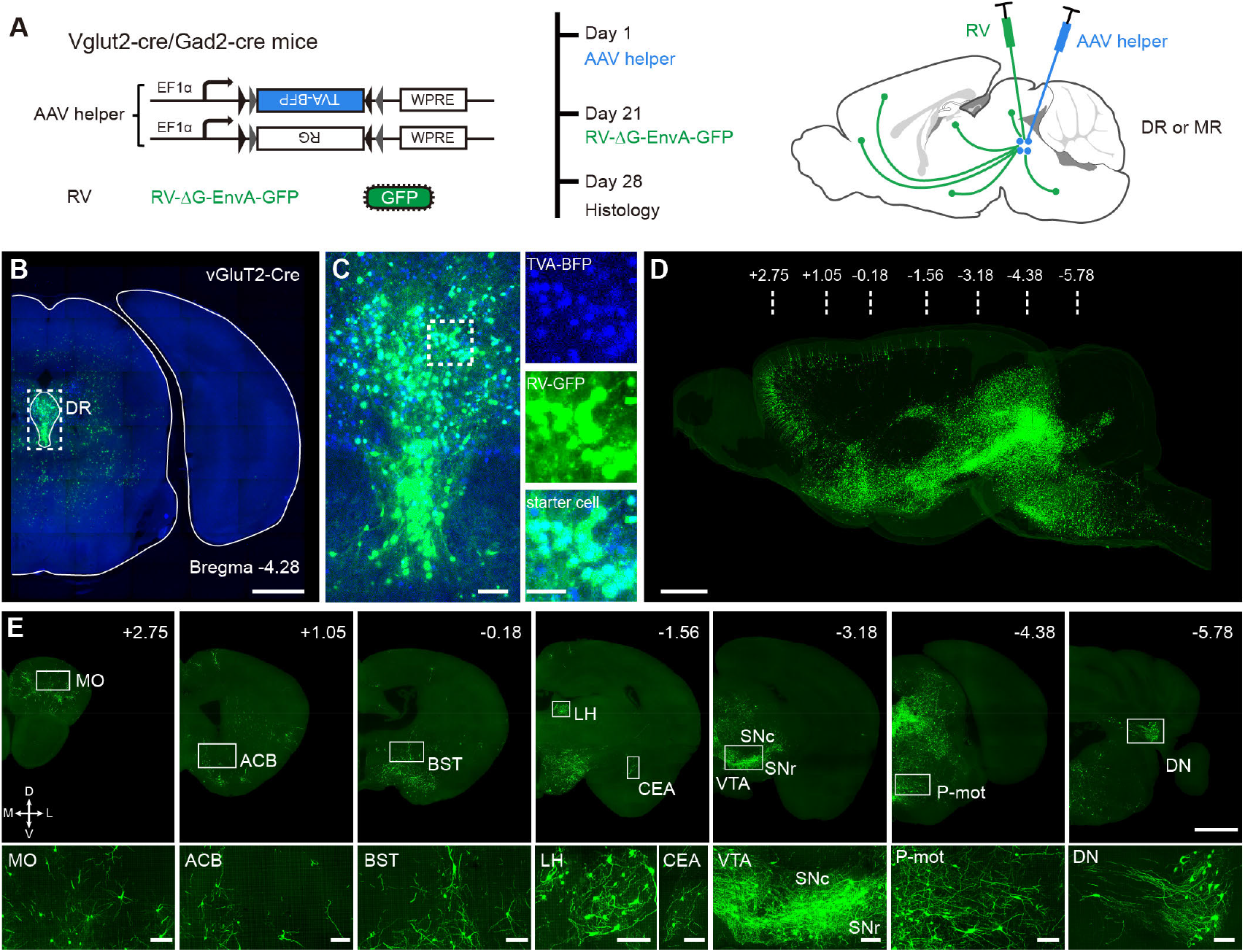
Whole-brain mapping of monosynaptic input neurons to cell-type-specific neurons in the DR and MR. (**A**) Schematic of monosynaptic rabies virus tracing the inputs to cell-type-specific neurons. The AAV helper virus expresses a fusion of TVA receptor and BFP and RG, and the modified rabies virus pseudotyped with EnvA expresses GFP. The experimental strategy and time line are on the right. (**B**) Representative schematic coronal section of the injection site. Scale bar, 1mm. (**C**) Left, enlarged view of dotted box area in (**B**) showing the starter cells (cyan). Right, enlarged view of the dotted box area on the left. The neurons co-labeled by BFP and GFP were starter cells (cyan), and GFP-labeled neurons were input neurons (green). Scale bars, left, 100 μm, right, 50 μm. (**D**) Three-dimensional rendering of whole-brain input neurons to DR glutamatergic neurons from a representative sample. Scale bar, 1mm. (**E**) Representative coronal sections of maximum intensity projection showing the distribution of input neurons to DR glutamatergic neurons. The projections are 50 μm thick. Scale bars, top row, 1 mm, bottom row, 100 μm. A, anterior; P, posterior; M, medial; L, lateral. The details of abbreviations for brain regions see Supplementary File 1.

To acquire the whole-brain high-resolution dataset, the virus-labeled samples were embedded in glycol methacrylate (GMA) resin and imaged with our home-made fMOST system (Gong et al., 2016) at a resolution of 0.32 × 0.32 × 2 μm^3^ (**Figure 1D,E**). Such high-resolution images indicate that the soma and neurite of labeled input neurons are finely detailed. From anterior to posterior, we observed dense input neurons in the isocortex, striatum, pallidum, thalamus, hypothalamus, midbrain, pons, medulla and cerebellar nuclei (**Figure 1—figure supplement 1**). However, in the olfactory areas, cortical subplate, hippocampus and cerebellar cortex, there were either no or sparse input neurons (**Figure 1—figure supplement 1**).

### Quantified whole-brain inputs to glutamatergic and GABAergic neurons in the DR and MR

To quantify the distributions of monosynaptic input neurons in each brain region, the coordinates of the soma of input neurons were detected using the NeuroGPS algorithm (Quan et al., 2013) and manually checked. The soma of input neurons were registered to the Allen Mouse Brain Common Coordinate Framework version 3 (Allen CCFv3) (**Figiure 2 A,B; Materials and methods**) (Ni et al., 2020; Wang et al., 2020). Based on Allen CCFv3’s hierarchy of brain regions, we identified 71 brain regions that have close connections with DR and MR neurons for further analysis (**Materials and methods; Supplementary File 1**).

**Figure 2.**
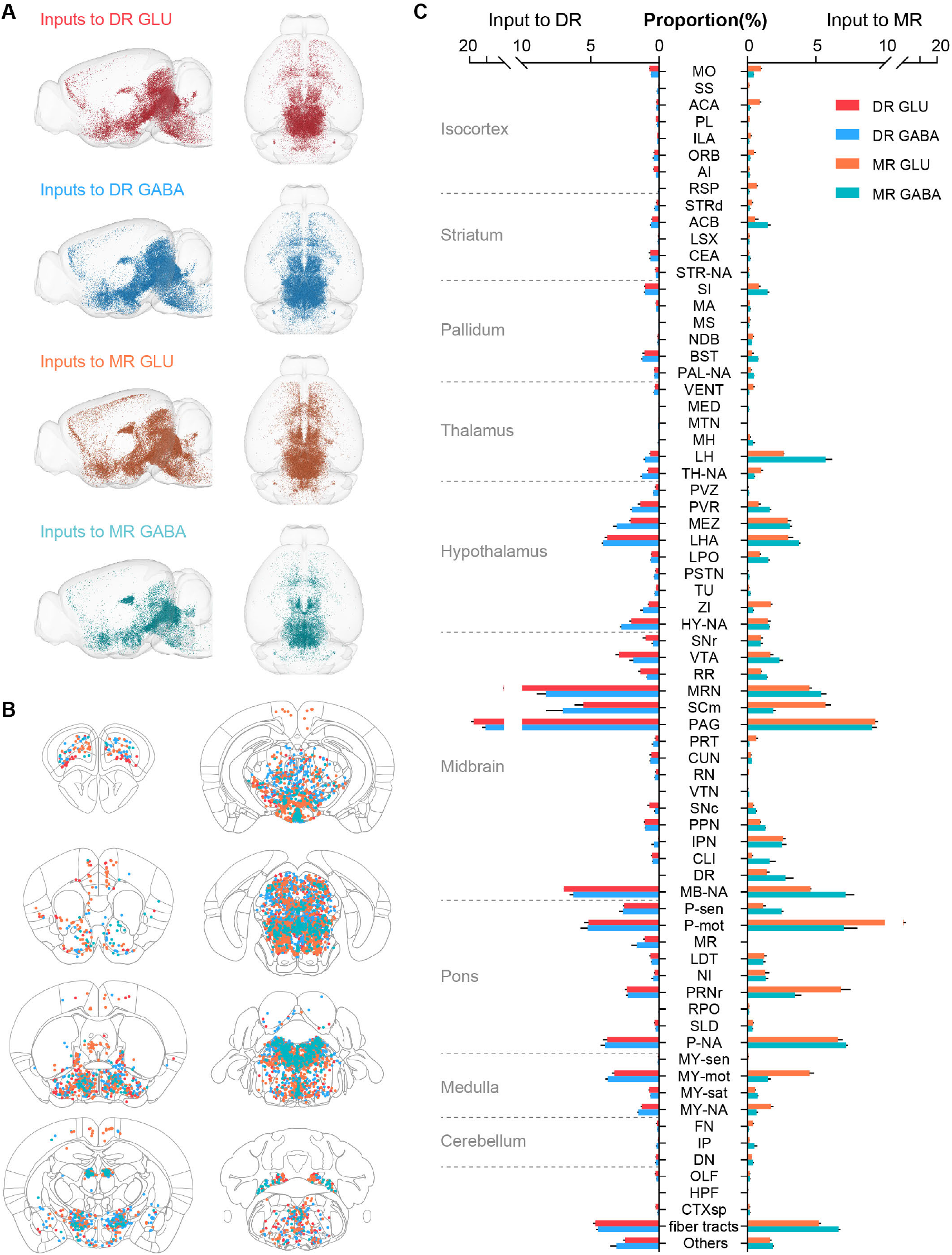
Whole-brain distribution of input neurons to glutamatergic and GABAergic neurons in the DR and MR. (**A**) Three-dimensional visualization of whole-brain inputs to glutamatergic neurons (GLU) and GABAergic neurons (GABA) in the DR and MR in representative samples. (**B**) Representative coronal sections illustrating the detected and registered input neurons. One dot represents one neuron, and the different colors reflect inputs to different types of neuron as in (**A**). Each section is 50 μm thick. (**C**) Proportion of the input neurons to glutamatergic and GABAergic neurons in the DR and MR across individual brain regions. Data are shown as mean ± s.e.m., n = 4 per group. The source data see Supplementary File 2. The details of abbreviations for brain regions see Supplementary File 1. The abbreviation NA indicates the non-annotated area in Allen CCFv3.

To generate the distribution of whole-brain input neurons, we calculated the number of input neurons in each brain region. To eliminate the variability in the total number of input neurons of different samples, the data were normalized by the total number of input neurons (excluding neurons in the target region) to get the proportion of input neurons in each brain region. Thus, we generated the quantified whole-brain distribution of long-range input neurons (**Figure 2C; Supplementary File 2**). To evaluate the consistency of the inputs to the same groups of neurons across different samples, we performed correlation analysis. The highly correlated results demonstrated the consistency and reliability of our data (**Figure 2—figure supplement 1 A,B**). We conducted unsupervised hierarchical clustering and bootstrapping of all samples. The input patterns of the four groups of neurons were divided into two clusters based on the target region, then input patterns of MR glutamatergic and GABAergic neurons were segregated based on neuron types (**Figure 2—figure supplement 1 C**).

### Comparison of inputs to glutamatergic and GABAergic neurons in the DR and MR

To explore the relationship of whole-brain long-range inputs to glutamatergic and GABAergic neurons in the DR and MR, first, we compared the inputs from the MR to DR glutamatergic and GABAergic neurons and found no significant difference (p=0.222, one-way ANOVA), then we compared inputs from the DR to MR glutamatergic and GABAergic neurons and also found no significant difference (p=0.069, one-way ANOVA). Next, we compared the whole-brain inputs to glutamatergic and GABAergic neurons in the DR and MR across brain regions using correlation analysis and variance analysis (one-way ANOVA followed by multiple comparisons with Tukey’s test; **Supplementary File 2**) (Ogawa et al., 2014). The whole-brain inputs to glutamatergic and GABAergic neurons in the same raphe nucleus were highly similar, while the whole-brain inputs to the same type of neurons in the DR and MR were similar with relatively lower correlation coefficients (**Figure 3A-D**). Whereas, there were quantitative differences in certain brain regions embedded in the overall similarity of the input patterns (**Figure 3A-D**).

**Figure 3.**
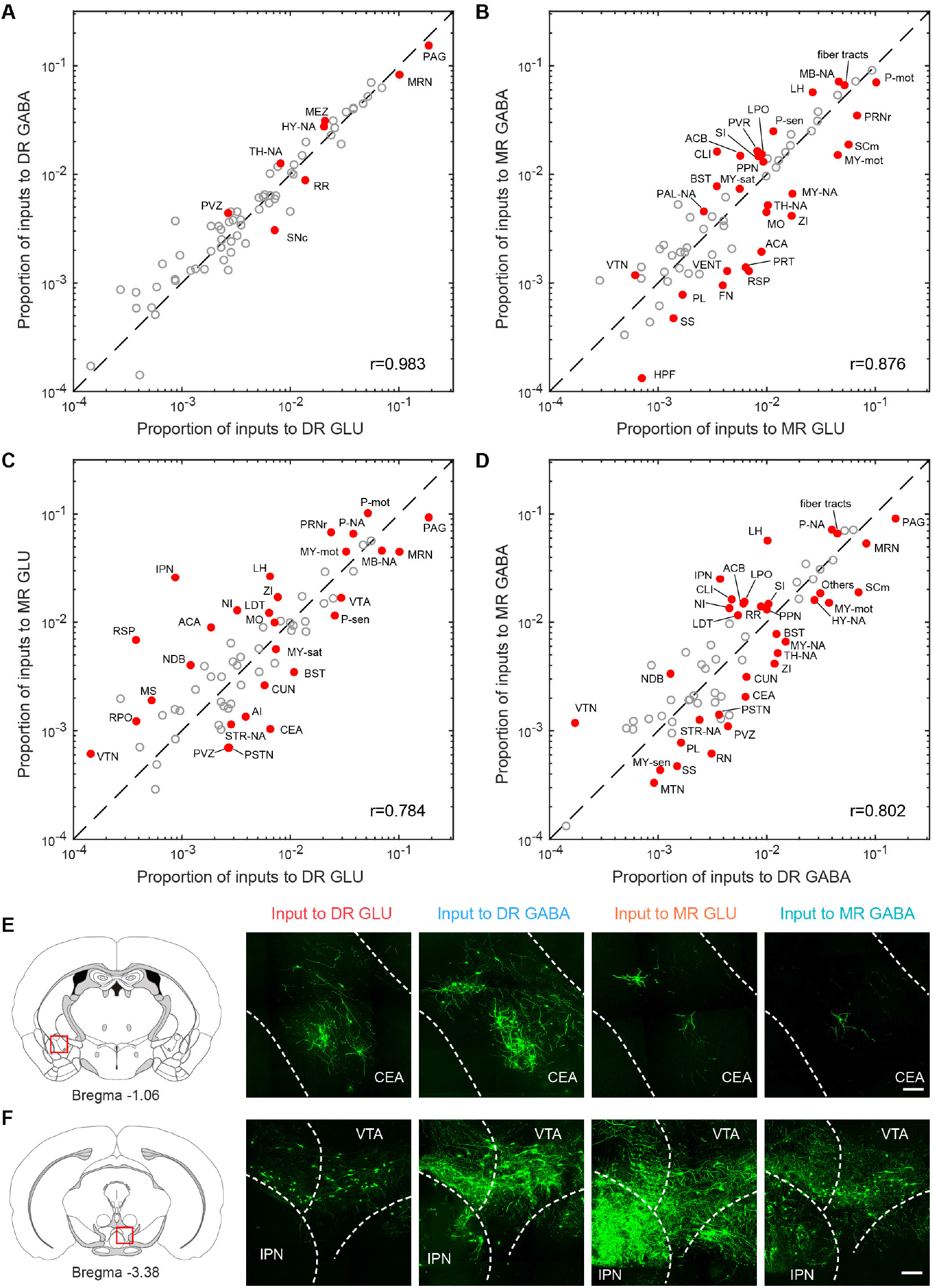
Comparisons of inputs to glutamatergic and GABAergic neurons in the DR and MR. (**A-D**) Comparisons of inputs to glutamatergic and GABAergic neurons in the DR and MR. The values for each region are the means of the proportion of input neurons from different samples of the same type. Red indicates significant differences (p< 0.05, One-way ANOVA followed by multiple comparisons with Tukey’s test). The p-values see Supplementary File 2. r: Pearson’s correlation coefficients. The details of abbreviations for brain regions see Supplementary File 1. (**E**) Comparison of input neurons in the CEA. Left: position of the images on the right. Right: RV-GFP-labeled input neurons in the CEA. Representative images are from maximum intensity projections of the coronal sections. The projections were 50 μm thick. Scale bar, 100 μm. (**F**) Comparison of input neurons in the IPN. Left: position of the images on the right. Right: RV-GFP-labeled input neurons in the IPN. Representative images are from maximum intensity projections of the coronal sections. The projections were 50 μm thick. Scale bar, 100 μm.

Specifically, a modest proportion of input neurons were distributed in the isocortex (**Figure 2C**). Several brain regions had biased inputs to different raphe neuron groups, especially the somatomotor areas (MO), anterior cingulate area (ACA) and retrosplenial area (RSP), which preferentially innervated MR glutamatergic neurons in comparison with MR GABAergic neurons and DR glutamatergic neurons (**Figure 3A-D**).

The striatum and pallidum contributed substantial inputs to glutamatergic and GABAergic neurons in the DR and MR (**Figure 2C**). Notably, the central amygdalar nucleus (CEA) preferentially innervated glutamatergic and GABAergic neurons in the DR compared with the MR (**Figure 3C-E**). And the BST sent prominent inputs to both glutamatergic and GABAergic neurons in the DR and MR with a preference for the DR (**Figure 3C,D**). Compared with DR neurons, MR neurons received a larger proportion of inputs from the diagonal band nucleus (NDB) (**Figure 3C,D**).

The main thalamic input neurons were assembled in the LH (**Figure 2C**). And there was a preference for LH neurons to have more inputs to MR glutamatergic and GABAergic neurons than to DR neurons (**Figure 3C,D**). Notably, MR GABAergic neurons received more inputs from the LH than MR glutamatergic neurons (**Figure 3B**). We also observed vast inputs from the hypothalamus to DR and MR glutamatergic and GABAergic neurons, in which the LHA provided the largest proportion, followed by the hypothalamic medial zone (MEZ), periventricular region (PVR), zona incerta (ZI) and lateral preoptic area (LPO) (**Figure 2C**). The ZI provided more inputs to MR glutamatergic neurons than to MR GABAergic neurons and DR glutamatergic neurons (**Figure 3B,C**). The LPO preferentially innervated MR GABAergic neurons in comparison with DR GABAergic neurons and MR glutamatergic neurons (**Figure 3B,D**).

The largest proportion of inputs to glutamatergic and GABAergic neurons in the DR and MR were from the midbrain (**Figure 2C**). Although glutamatergic and GABAergic neurons in the DR and MR received massive inputs from the periaqueductal gray (PAG) and midbrain reticular nucleus (MRN), DR neurons received more than MR neurons (**Figure 3C,D**). The interpeduncular nucleus (IPN) provided remarkable inputs to glutamatergic and GABAergic neurons in the MR with very sparse inputs to the DR (**Figure 3C,D,F**). The superior colliculus, motor related (SCm) contributed more inputs to MR glutamatergic neurons and DR GABAergic neurons than to MR GABAergic neurons (**Figure 3B,D**).

The pons contributed dense inputs, in which the pons, motor related (P-mot) and pontine reticular nucleus (PRNr) preferentially innervated MR glutamatergic neurons in comparison with MR GABAergic neurons and DR glutamatergic neurons (**Figure 3B,C**). However, the pons, sensory related (P-sen) provided more inputs to DR glutamatergic neurons than to MR glutamatergic neurons (**Figure 3C**). Moreover, the medulla, motor related (MY-mot) preferentially provided inputs to MR glutamatergic neurons in comparison with MR GABAergic neurons and DR glutamatergic neurons (**Figure 3B,C**). These results indicated that the glutamatergic and GABAergic neurons in the raphe nucleus receive inputs from similar upstream brain regions with quantitative differences in certain brain regions.

### Whole-brain outputs of glutamatergic and GABAergic neurons in the DR and MR

To systematically map whole-brain outputs of glutamatergic and GABAergic neurons in the DR and MR, we stereotaxically injected Cre-dependent AAV-DIO-EYFP into the DR or MR in Vglut2-Cre and Gad2-Cre mice (n=4 per group). The virus-labeled and GMA resin-embedded samples were imaged using fMOST system (**Figure 4A**). To generate quantified outputs of glutamatergic and GABAergic neurons in the DR and MR, we registered the high-resolution whole-brain image datasets to Allen CCFv3 and segmented the projection signal to calculate the proportion of projection signal across brain regions (**Figure 4A,B; Supplementary File 3; Materials and methods**).

**Figure 4.**
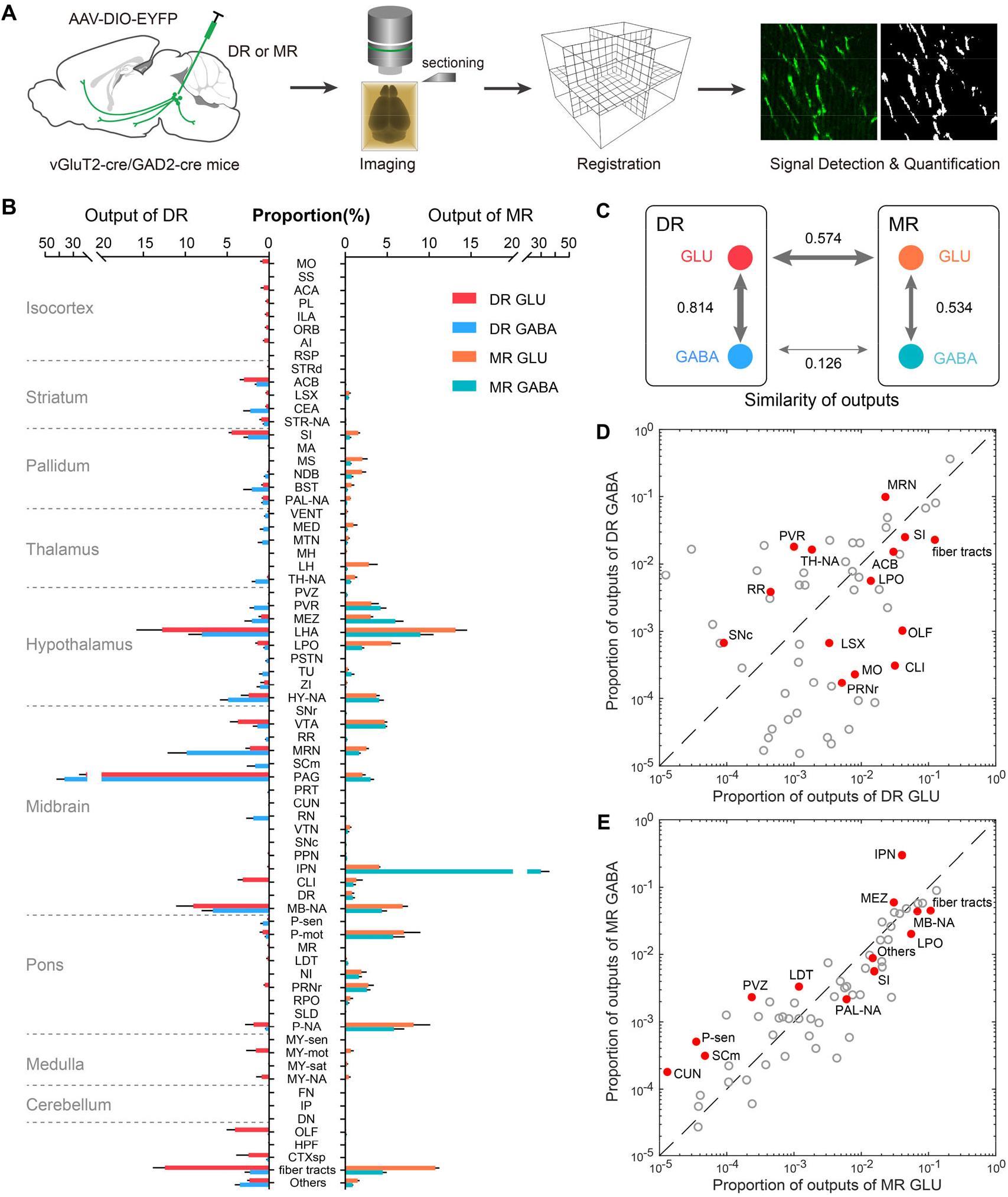
Whole-brain outputs of glutamatergic and GABAergic neurons in the DR and MR. (**A**) Schematic outlining viral tracing, whole-brain imaging, data processing and analysis. (**B**) Proportion of the outputs of glutamatergic and GABAergic neurons in DR and MR across individual brain regions. Data are shown as mean ± s.e.m., n=4 per group. The source data see Supplementary File 3. (**C**) Similarities of whole-brain projection patterns. The numbers indicate Pearson’s correlation coefficients. The arrow thickness indicates the magnitude of similarity. (**D**) Comparison of outputs of glutamatergic and GABAergic neurons in the DR. (**E**) Comparison of outputs of glutamatergic and GABAergic neurons in the MR. The values for each region are the means of the proportion of outputs from different samples of the same type. Red indicates significant differences (p< 0.05, one-way ANOVA). The p-values see Supplementary File 3. The details of abbreviations for brain regions see Supplementary File 1.

On the whole-brain level, glutamatergic and GABAergic neurons in the DR and MR provided substantial ascending projections to the forebrain and midbrain and varying degrees of descending projections to the pons and medulla (**Figure 4B; Figure 4—figure supplement 1**). MR glutamatergic and GABAergic neurons predominately innervated midline structures, while DR glutamatergic and GABAergic neurons projected more broadly and laterally, and most of their projection targets were distinctive. Regarding the forebrain, DR neurons projected more broadly to the prefrontal cortex, amygdala, nucleus accumbens (ACB) and BST, while MR neurons innervated the lateral septal complex, medial septal nucleus (MS), NDB and LH (**Figure 4B; Figure 4—figure supplement 1**). And MR glutamatergic and GABAergic neurons had more outputs to the pons in comparison with DR neurons (**Figure 4B**). Meanwhile, they both sent dense projections to the hypothalamus and midbrain areas, such as the LHA and ventral tegmental area (VTA) (**Figure 4B; Figure 4—figure supplement 1**). Moreover, glutamatergic neurons seemed to project more broadly than GABAergic neurons, while GABAergic neurons preferentially innervated neighboring brain regions, such as the PAG and MRN for DR GABAergic neurons and the IPN for MR GABAergic neurons.

We quantitatively compared the projection patterns of glutamatergic and GABAergic neurons in the DR and MR. The same types of neurons in the DR and MR have divergent projection patterns (**Figure 4C**). Regarding the glutamatergic and GABAergic neurons in the same raphe nucleus, although their overall projection patterns were relatively similar, there were differences in critical brain regions (**Figure 4C-E; Supplementary File 3**). Notably, regarding the amygdala, DR GABAergic neurons mainly projected to the CEA with scarce projections to the basolateral amygdalar nucleus (BLA), while DR glutamatergic neurons preferentially projected to the BLA (**Figure 4—figure supplement 2A,D**). And DR GABAergic neurons sent considerable projections to the DMH and paraventricular nucleus of the thalamus (PVT), while there were scarce or no axonal projections of DR glutamatergic neurons in these regions (**Figure 4—figure supplement 2B-D**). Regarding MR neurons, we found dense projections of glutamatergic neurons in the LH but scarce projections of GABAergic neurons (**Figure 4B**). And the IPN received 29.9% of the total projections from MR GABAergic neurons but only 4.0% from MR glutamatergic neurons (**Figure 4B,E**).

### Habenula-raphe circuits

The habenula, consisting of the medial habenula (MH) and lateral habenula (LH), appears to be a node connecting the forebrain and midbrain regions that are related to emotional behaviors (Hikosaka, 2010). The LH has been closely connected to the DR and MR both anatomically and functionally, and their connections are involved in aversion-related behavior and depression (Hu et al., 2020; Zhao et al., 2015). The LH provided dense inputs to glutamatergic and GABAergic neurons in the DR and MR with a preference to MR neurons in comparison with corresponding DR neurons (**Figure 3C, D; Figure 5A**). And the input neurons to MR glutamatergic and GABAergic neurons were assembled more caudally (**Figure 5A**). Specifically, as MR GABAergic neurons received more inputs from the LH than MR glutamatergic neurons (**Figure 3D**), we found that the lateral part of LH sent dense inputs to MR GABAergic neurons but sparser inputs to MR glutamatergic neurons and that the input neurons to MR GABAergic neurons were assembled more laterally than the input neurons to MR glutamatergic neurons on the whole (**Figure 5A**).

**Figure 5.**
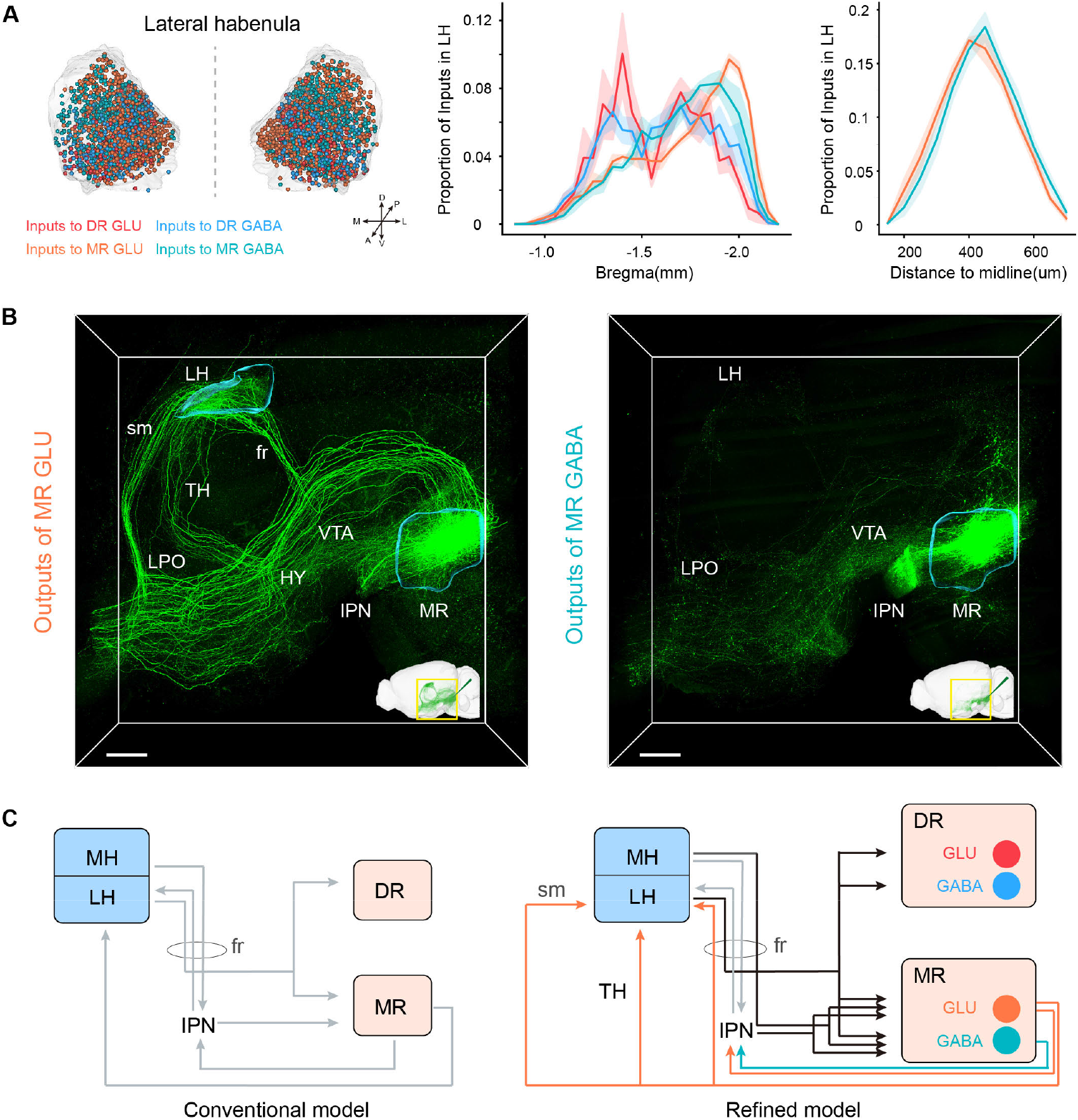
Habenula-raphe Circuit. (**A**) Comparison of inputs in the LH to glutamatergic and GABAergic neurons in the DR and MR. Left, three-dimensional rendering of input neurons in the LH from representative samples. One dot represents one neuron, and the different colors reflect inputs to different types of neurons. Middle, density plot of the proportion of input neurons in the LH along the anterior-posterior axis. Right, density plot of the proportion of input neurons to MR glutamatergic and GABAergic neurons in the LH along the medial–lateral axis. Bin width, 50 μm. The shaded area indicates s.e.m., n=4. (**B**) Representative projections of MR glutamatergic and GABAergic neurons. The image is a perspective view of three-dimensional rending of projections in the region of interest shown in the bottom right corner. The image in the bottom right corner is three-dimensional rending of projections in the left hemisphere. The rendered data have been registered to Allen CCFv3. Scale bar, 500 μm. (**C**) A refined model of the habenula-raphe circuit based on connections with glutamatergic and GABAergic neurons in the DR and MR. The conventional model is from previous studies (Hikosaka, 2010; Hu et al., 2020). In the refined model, the inputs identified in this study are shown in black, the outputs of MR glutamatergic and GABAergic neurons are shown in orange and turquoise respectively, and the known circuits are shown in gray. TH, thalamus; HY, hypothalamus; sm, stria medullaris; fr, fasciculus retroflexus.

However, there were no projections from DR glutamatergic and GABAergic neurons and scarce projections from MR GABAergic neurons to the LH, only MR glutamatergic neurons sent strong projections to the LH (mainly assembled in the medial part of LH) (**Figure 4B**; **Figure 5—figure supplement 1A,C**). Taking advantage of our three-dimensional high-resolution imaging, we found that MR Vglut2+ neurons send projections to the LH through multiple pathways: through the fasciculus retroflexus, stria medullaris and thalamus respectively (**Figure 5B; Figure 5— figure supplement 1B,C**). There was evidence that Vglut2+ neurons in surrounding regions of the MR do not project to the LH (Szőnyi et al., 2019), which supported the reliability of this projection pattern. The specific reciprocal connections between MR glutamatergic neurons and the LH suggested that MR glutamatergic neurons might be involved in some specific functions related to the LH. The LH has been revealed to play a critical role in aversion and depression (Cui et al., 2018; Hu et al., 2020; Yang et al., 2018), and previous studies discovered that MR Vglut2+ neurons could activate the LH and that activation of MR Vglut2+ neurons could induce aversive behaviors and depressive symptoms (Szőnyi et al., 2019). These results highlight the importance of the structural characteristics of the MR-LH pathway for their function roles.

Although the MH sent few inputs to MR glutamatergic and GABAergic neurons, and scarce inputs to the DR, it has been thought to strongly project to the IPN (Lima, et al., 2017; Qin et al., 2009). In our results, the IPN had remarkable inputs to MR glutamatergic and GABAergic neurons with very sparse inputs to DR glutamatergic and GABAergic neurons (**Figure 3F**). These results implied that MR glutamatergic and GABAergic neurons receive inputs directly and indirectly (via the IPN) from the MH. Moreover, MR glutamatergic and GABAergic neurons strongly projected to the IPN (**Figure 5B**), and the IPN has been revealed to project to the LH (Lima, et al., 2017). These results indicated the sophisticated connections of the habenula, IPN, DR and MR. Based on the conventional model of the habenula-raphe circuit (Hikosaka, 2010; Hu et al., 2020), we proposed a more refined model of the habenula-raphe circuit (**Figure 5C**).

### Whole-brain connectivity pattern of glutamatergic and GABAergic neurons in the DR and MR

The glutamatergic and GABAergic neurons in the DR and MR received inputs from and sent outputs to a wide range of brain regions (**Figure 6—figure supplement 1A**). We assessed the similarities between the whole-brain inputs and outputs of the same group of neurons using Pearson’s correlation coefficient. Regarding glutamatergic and GABAergic neurons in the DR, the correlation coefficients were 0.766 and 0.839 respectively (**Figure 6—figure supplement 1A**).

These results indicated that they have reciprocal connections with vast brain regions, which were mainly in the striatum, pallidum, hypothalamus and midbrain (**Figure 2C; Figure 4B**). Regarding glutamatergic and GABAergic neurons in the MR, the correlation coefficients were 0.578 and 0.384 respectively (**Figure 6—figure supplement 1A**), which was related to the fact that MR neurons receive massive inputs from the isocortex, striatum and medulla but sparsely project to these regions (**Figure 2C; Figure 4B**).

There were massive reciprocal connections of glutamatergic and GABAergic neurons in the DR and MR, which implied the feedback regulation of specific functions. To assess the reciprocity, we calculated the ratio of the proportion of outputs to the proportion of inputs for each brain region (**Figure 6—figure supplement 1B**). Approximately 45% brain regions had ratio value between 0.25 and 4, indicating relatively balanced reciprocal connectivity. In contrast, approximately 45% brain region had input bias (ratio value<0.25), and only a few brain regions showed output bias (ratio value>4). Particularly, for MR GABAergic neurons, the IPN contributed 2.5% of all inputs but received 29.9% of outputs.

The glutamatergic and GABAergic neurons in the DR and MR receive a vast range of inputs, whereas the relationship between different upstream brain regions needs to be explored. And the axons of neurons have collateral branches targeting different areas, but the relationship of these targets also needs further exploration. Based on the fact that DR and MR were heterogeneous and each injection only labeled a part of neurons that might have different connectivity, we performed correlation analysis and hierarchical cluster analysis to explore the similarities and variances of inputs and outputs of brain regions connected with glutamatergic and GABAergic neurons in the DR and MR. We selected 14 brain regions that have close long-range connections with glutamatergic and GABAergic neurons in the DR and MR for analysis. As a result, the clusters were not completely consistent regarding the inputs or outputs of glutamatergic and GABAergic neurons in a particular raphe nucleus (**Figure 6A,B**). Regarding inputs to DR glutamatergic and GABAergic neurons, upstream brain regions formed clear clusters, but the clusters were not consistent, where only the substantia nigra, reticular part (SNr), substantia nigra, compact part (SNc) and VTA formed the same cluster. Regarding outputs of DR glutamatergic neurons, the MEZ, LPO, LHA, ZI, SNr and IPN formed a cluster that had very high correlation. In contrast, for outputs of DR GABAergic neurons, different clusters were presented. These results implied that DR glutamatergic and GABAergic neurons might have different collateral projection patterns. In addition, a pair of brain regions might display opposing correlations regarding the inputs or outputs of glutamatergic and GABAergic neurons in the DR. For instance, the substantia innominate (SI) and NDB were positively correlated for inputs to DR glutamatergic neurons but negatively correlated for inputs to DR GABAergic neurons (**Figure 6A**), which suggested that DR glutamatergic and GABAergic neurons might receive distinct inputs from basal forebrain.

**Figure 6.**
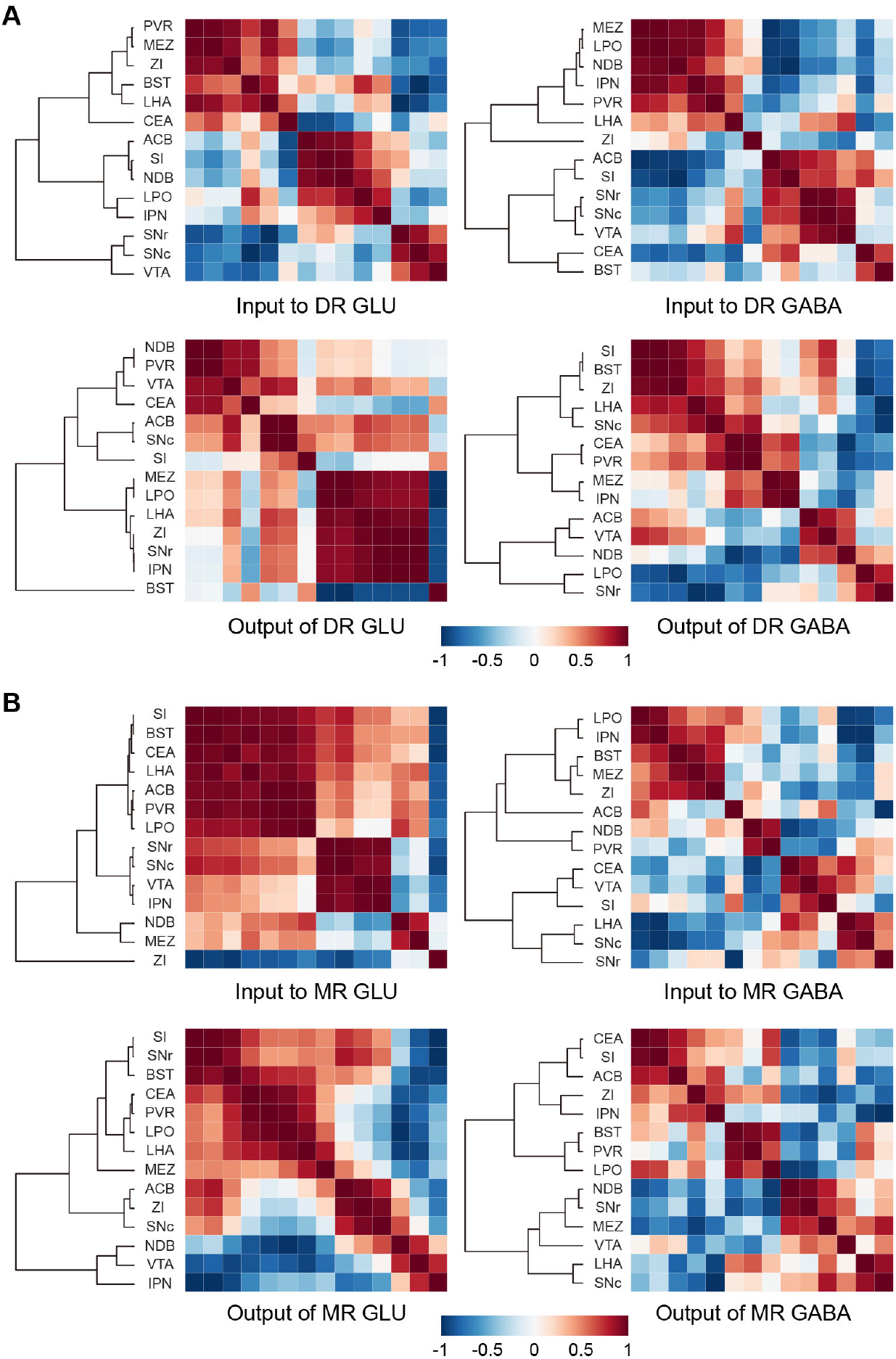
Connectivity patterns of glutamatergic and GABAergic neurons in the DR and MR. (**A**) Correlation and hierarchical cluster analysis showing the similarities and variances in brain regions connected with DR glutamatergic and GABAergic neurons. The heatmap represents Pearson’s correlation coefficient matrix. (**B**) Correlation and hierarchical cluster analysis showing the similarities and variances in brain regions connected with MR glutamatergic and GABAergic neurons. The heatmap represents Pearson’s correlation coefficient matrix. The details of abbreviations for brain regions see Supplementary File 1.

Regarding inputs and outputs of MR glutamatergic neurons, clear clusters formed. Specifically, as for the clusters of inputs to MR glutamatergic neurons, the ZI was separate from other regions, which indicated the ZI-MR glutamatergic neurons pathway might execute special functions different from other upstream brain regions. Regarding inputs and outputs of MR GABAergic neurons, no obvious clusters formed, which implied that MR GABAergic neurons might have few collateral projections to these brain regions. And there were also pairs of brain regions displaying opposing correlations regarding the inputs or outputs of MR glutamatergic and GABAergic neurons, such as the SNc and IPN were positively correlated for inputs to MR glutamatergic neurons but negatively correlated for inputs to MR GABAergic neurons (**Figure 6B**). Overall, these results implied that glutamatergic and GABAergic neurons in the DR and MR might receive inputs from and project to various unions of brain regions.

## Discussion

In this study, we used virus tracing and whole-brain high-resolution imaging to generate a comprehensive whole-brain atlas of inputs and outputs of glutamatergic and GABAergic neurons in the DR and MR (**Figure 7A,B**). And systematic quantitative analysis has been performed to elucidate the convergences and divergences in input and output patterns.

**Figure 7.**
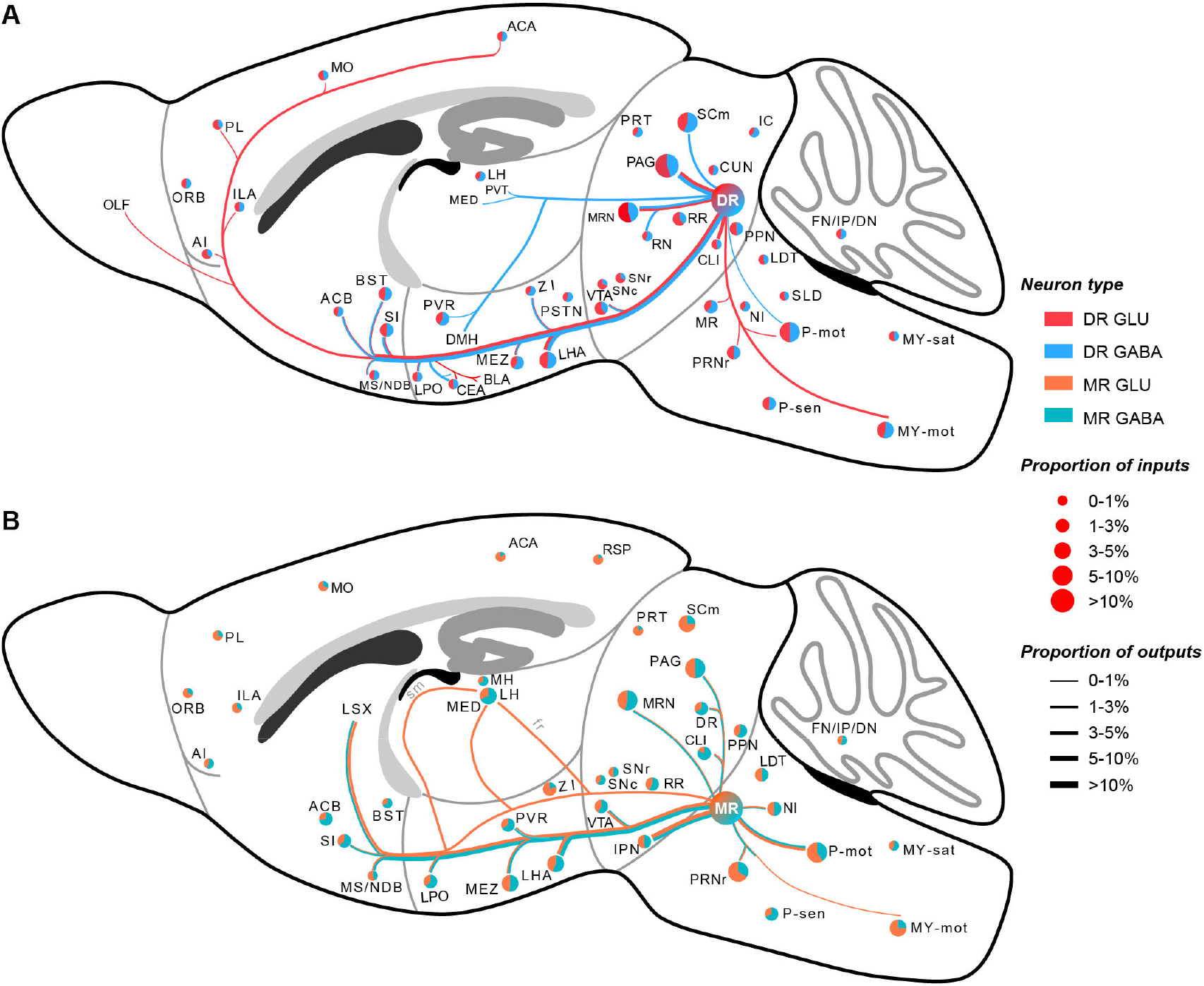
Whole-brain schematic of the inputs and outputs of glutamatergic and GABAergic neurons in the DR and MR. (**A**) Whole-brain schematic of the inputs and outputs of glutamatergic and GABAergic neurons in the DR. (**B**) Whole-brain schematic of the inputs and outputs of glutamatergic and GABAergic neurons in the MR. The pie charts represent the inputs in each brain region, where colors reflect the postsynaptic neuron types, and the size reflects the proportion value. The lines represent outputs in each brain region, where colors reflect the neuron types, and the line thickness reflects the proportion value. The details of abbreviations for brain regions see Supplementary File 1.

We found that glutamatergic and GABAergic neurons in the DR and MR receive inputs from similar upstream brain regions but send distinctive outputs. There were essential differences between the connections of DR and MR neurons. DR glutamatergic and GABAergic neurons had close connections with the CEA, while MR glutamatergic and GABAergic neurons received few inputs from CEA and did not project to the CEA (**Figure 6—figure supplement 1A**). And the CEA has been revealed to regulate reward and food intake (Carter, et al., 2013; Janak et al., 2015; Zséli et al., 2018) that are also regulated by DR glutamatergic and GABAergic neurons (Nectow et al., 2017). The IPN had dense reciprocal connections with MR glutamatergic and GABAergic neurons, but almost no direct connections with DR glutamatergic and GABAergic neurons (**Figure 6—figure supplement 1A**). The IPN and MR are both considered as important parts of the midline network involved in regulating hippocampal theta rhythm (Lima et al., 2017). The consistency between anatomical connectivity and behavioral function indicated the significance of dissecting whole-brain connectivity for elucidating functions.

The differences between connections of glutamatergic and GABAergic neurons in the same raphe nucleus might provide insight into their functions. DR GABAergic neurons preferentially projected to the CEA, unlike DR glutamatergic neurons (**Figure 4—figure supplement 1A,D**). Optogenetic activation of the CEA inhibits food intake (Carter et al., 2015), while activation of DR GABAergic neurons increases food intake (Nectow et al., 2017). Furthermore, DR GABAergic neurons uniquely innervate the PVT (**Figure 4—figure supplement 1C,D**), which is connected with the CEA and involved in inhibiting food intake (Kirouac, 2015). Thus, we speculated that activation of DR GABAergic neurons might inhibit CEA and PVT neurons to increase food intake. DR GABAergic neurons send considerable projections to DMH, while DR glutamatergic neurons scarcely project to the DMH (**Figure 4—figure supplement 1B,D**). And DR GABAergic neurons have been revealed to regulate thermogenesis via projections to the DMH (Schneeberger et al., 2019). These results indicated that DR GABAergic neurons might play a more critical role in regulating thermogenesis than DR glutamatergic neurons. We found that MR glutamatergic neurons project to the LH via multiple pathways while MR GABAergic neurons scarcely project to the LH (**Figure 5B**), indicating that there might be different subtypes of MR glutamatergic neurons projecting to LH via different pathways. There is evidence that MR Vglut2+ neurons could activate the LH (related to aversion and negative prediction) and control the acquisition of negative experience (Szőnyi et al., 2019). Our results implied that MR glutamatergic neurons that project to the LH via multiple pathways might regulate the different aspects of aversive and negative emotions. These results and implications highlight the biased or unique connectivity of different types of neurons in the same raphe nucleus are related to the regulation of specific functions. Our work could provide the resource for dissecting the functions of glutamatergic and GABAergic neurons in the DR and MR.

It should be noted that a pair of brain regions might display different or even opposing correlations regarding the inputs to glutamatergic and GABAergic neurons in the same raphe nucleus **(Figure 6 A,B**), which implied that different neuron groups in upstream brain regions might individually target heterogeneous raphe neuron groups. More advanced labeling methods that could label upstream inputs to different cell types in a single brain sample is needed to explore this problem further. The target regions of projections of glutamatergic and GABAergic neurons in the same raphe nucleus also showed different correlations (**Figure 6 A,B**). This result indicated that glutamatergic and GABAergic neurons in the same raphe nucleus might have different collateral projection patterns, which are worth illustrating through complete single neuron reconstruction.

Our results were in accordance with the input and output circuits of neurons in DR and MR identified using classic tracing techniques (Marcinkiewicz et al., 1989; Oh, et al., 2014; Peyron et al., 1997; Vertes et al., 2008). However, the DR and MR are heterogeneous and contain diverse types of neurons, including glutamatergic, GABAergic, serotonergic and dopaminergic neurons. In comparison with the known circuits of specific types of neurons in the DR and MR, our results matched well with the current incomplete knowledge of input and output circuits of glutamatergic and GABAergic neurons in the DR and MR, and we found that different neuron groups in the same raphe nucleus received inputs from similar upstream brain regions and sent complementary projections (Lin et al., 2020; Muzerelle et al., 2016; Ogawa et al., 2014; Pollak Dorocic et al., 2014; Ren et al., 2018a; Ren et al., 2019; Weissbourd et al., 2014). There are essential brain regions having biased connections with different types of neurons in the same raphe nucleus. In the DR, serotonergic neurons project more broadly, such as to the entorhinal area and piriform area (Ren et al., 2018a; Ren et al., 2019). Moreover, a key difference is that DR serotonergic neurons send projections to the lateral habebula (Muzerelle et al., 2016; Ren et al., 2018a; Zhang et al.,2018), whereas DR glutamatergic and GABAergic neurons do not (**Figure 4B**). In the MR, previous studies have revealed that MR neurons project to the hippocampus and regulate multiple hippocampal activities (Jackson et al., 2008; Varga et al., 2009; Vertes et al., 2008). In this study, we did not observe apparent projections of MR Vglut2+ and Gad2+ neurons in the hippocampus (**Figure 4B**), which was in accordance with the previous studies that most retrogradely labeled MR neurons to the hippocampus are serotonergic or Vglut3-positive (Szőnyi et al., 2016). The biased connections caused by cell-type specificity emphasize the necessity of dissecting the whole-brain connectivity of different cell types in the same region.

From previous studies, GABAergic neurons in the DR and MR were thought to innervate other cell types in the raphe nucleus to modulate function through disynaptic pathways. Suppression of DR GABAergic neurons could alleviate the acquisition of social avoidance by promoting the activity of serotonergic neurons (Challis et al., 2013). MR GABAergic neurons modulate hippocampal theta rhythm by innervating MR serotonergic neurons, and indirectly regulate hippocampal ripple activity by inhibiting MR non-GABAergic neurons (Li, et al., 2005; Wang et al., 2015). However, DR GABAergic neurons also regulate thermogenesis through long-range projections to the DMH, BST and related areas (Schneeberger et al., 2019). As we found a vast range of projections from GABAergic neurons in DR and MR, it indicated an underappreciated potential functional role of GABAergic projection neurons in the DR and MR. Whether the same GABAergic neurons in the raphe nucleus could participate in the direct and indirect pathways simultaneously needs further investigation.

In summary, we constructed a comprehensive whole-brain atlas of inputs and outputs of glutamatergic and GABAergic neurons in the DR and MR, revealing similar input patterns but divergent projection patterns. The differences in connectivity patterns are related to specific regulatory processes of specific functions. As the whole-brain connections of genetically targeted neurons are key factors in characterizing cell types, our result would contribute to generating whole-brain cell atlases that are under ongoing effort. Our work could form the foundation for exploring the relationship of cell heterogeneity, anatomical connectivity and behavior function of the raphe nucleus.

## Materials and Methods

### Animals

In this study, adult Vglut2-Cre mice (stock number: 016963) and Gad2-Cre mice (stock number: 010802) purchased from the Jackson Laboratory were used. All mice were housed in an experiment environment with 12-h light/dark cycle, 22 ± 1 °C temperature, 55 ± 5% humidity and food and water ad libitum. All animal experiments were approved by the Institutional Animal Ethics Committee of Huazhong University of Science and Technology and were conducted in accordance with relevant guidelines.

### Stereotaxic injections

For retrograde monosynaptic tracing, 150 nl adeno-associated helper virus (AAV helper) was injected into the DR (bregma: −4.6 mm, lateral: 0 mm, ventral: −3.0 mm) or MR (bregma: −4.6 mm, lateral: 0 mm, ventral: −4.25 mm) in Vglut2-Cre and Gad2-Cre mice. Three weeks later, 200 nl RV-ΔG-EnvA-GFP (2×10^8^ infectious units/ml) was injected into the same site. One week later, the mice were used for sample preparation. The AAV helper was a 1:2 mixture of rAAV2/9-EF1α-DIO-His-TVA-BFP (2×10^12^ viral genomes/ml) and rAAV2/9-EF1α-DIO-RG (2×10^12^ viral genomes/ml). For antegrade tracing, 50 nl AAV-DIO-EYFP (2×10^12^ viral genomes/ml) was injected into the DR (bregma: −4.6 mm, lateral: 0 mm, ventral: −3.0 mm) or MR (bregma: −4.6 mm, lateral: 0 mm, ventral: −4.25 mm) in adult Vglut2-Cre and Gad2-Cre mice. Three weeks later, the mice were used for sample preparation. All viral tools were produced by BrainVTA Co., Ltd.

### Histology

All histological procedures followed a previously described workflow (Ren et al., 2018b). Briefly, the anesthetized mice were intracardially perfused with 0.01 M PBS (Sigma-Aldrich Inc), followed by 4% paraformaldehyde (Sigma-Aldrich Inc) in 0.01 M PBS. The brains were excised and post-fixed in 4% paraformaldehyde at 4 °C for 24 h. Subsequently, each brain was rinsed overnight at 4 °C in 0.01 M PBS and dehydrated in a graded ethanol series (50, 70 and 95% ethanol, changing from one concentration to the next every 1 h at 4 °C). After dehydration, the brains were immersed in a graded glycol methacrylate (GMA) series (Ted Pella Inc.), including 0.2% SBB (70%, 85%, and 100% GMA for 2 h each and 100% GMA overnight at 4 °C). Finally, the samples were impregnated in a prepolymerization GMA solution for 3 days at 4 °C and embedded in a vacuum oven at 35 °C for 24 h.

### Imaging and image preprocessing

For whole-brain high-resolution imaging, the virus-labeled and GMA resin embedded samples were imaged with propidium iodide (PI) simultaneously staining cytoarchitecture landmarks using our home-made fMOST system at a resolution of 0.32 μm× 0.32 μm × 2 μm. The acquired two-channel raw data are processed by mosaic stitching and illumination correction to piece together into entire coronal sections using the a previously described workflow (Gong et al., 2016). Each channel dataset of one brain sample contains approximately 5,500 coronal slices. For starter cells, the samples were sectioned in 50 μm coronal slices using the vibrating slicer (VT1200S, Leica) and imaged using the automated slide scanner (VS120 Virtual Slide, Olympus).

### Data processing

#### Registration

To quantify and integrate the whole-brain connections, the coordinates of the soma of input neurons and high-resolution image stack of labeled outputs were registered to Allen CCFv3 using the transformation parameters acquired by the previously described methods (Ni et al., 2020). In brief, we segmented several brain regions as landmarks through cytoarchitecture references, such as the outline, caudoputamen, medial habenula, lateral ventricle, third ventricle. Based on these landmarks, we performed affine transformation and symmetric image normalization in Advanced Normalization Tools (ANTS) to acquire transformation parameters.

#### Nomenclature of brain regions

Demarcation and annotation of brain regions were based on Allen CCFv3. The superior central nucleus raphe (CS) corresponds to the median raphe nucleus (MR) with the consultation of the mouse brain atlas by Paxinos and Franklin (Paxinos and Franklin, 2012). Based on Allen CCFv3’s hierarchy of brain regions, as there are no or few input neurons and projections in many brain regions, we collapsed some brain regions to their “parent” region as needed, thereby divided the whole-brain into 117 brain regions (see Supplementary File 1) and identified 71 brain regions for analysis (areas that have small proportion of connections are merged into “Others”). The STR-NA, PAL-NA, TH-NA, HY-NA, MB-NA, P-NA and MY-NA respectively refer to the non-annotated area in the striatum, pallidum, thalamus, hypothalamus, midbrain, pons and medulla.

#### Detection and quantification of whole-brain inputs

Regarding the input circuits, we automatically identified and localized the soma of input neurons using NeuroGPS (Quan et al., 2013) and manually checked the results to eliminate some indiscernible mistakes, then warped the soma coordinates to Allen CCFv3 using the transformation parameters from registration described above. We calculated the number and proportion of input neurons in each brain region of interest (excluding target area) to generate the quantified whole-brain inputs.

#### Detection and quantification of whole-brain outputs

Regarding the output circuits, we generated quantified whole-brain outputs by taking following steps:

We resampled the image stack of labeled neural structures to isotropic 1 μm, segmented the outline of brain and set the intensity of pixels outside the outline to 0, then used the transformation parameters of registration described above to warp them to Allen CCFv3 at 1 μm scaling. Then we manually segmented injection sites on registered coronal sections.

To detect projection signal from background, each registered coronal section was background subtracted, Gaussian filtered, and threshold segmented to binary image. The background image *I*^*^ was calculated as *I*^*^ = min(*I*,*backgroun d*) followed by ten convolutions with the averaging template of 9×9 size, where *I* is the gray level of coronal section and the backgroud is an approximate estimated background intensity (Quan et al., 2013). The size of gaussian filter was 5×5. The filtered image was binarized by 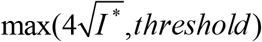 where *threshold* was the result calculated by the Yen method that clipped to the predetermined threshold range (Yen et al., 1995).

The whole-brain images were divided into 10×10×10 μm^3^ grids. In each division, we calculated signal density by the definition of the sum of detected pixels divided by the sum of all pixels in a three-dimensional grid, therefore generated a three-dimensional signal density matrix of 10 μm voxel resolution. Then we calculated the computational path based on the signal density matrix using multistencils fast marching algorithm and removed the voxels that could not back-track to injection site or back-track to injection site with low confidence (Oh et al., 2014; Hassouna and Farag, 2007; Liu, et al., 2018). The confidence of the path was defined as the proportion of back-tracking points that were located in the foreground voxel in the path, and the foreground voxel refers to the voxel whose signal density was greater than a threshold. Finally, we manually inspected the results and removed the remaining confusing noise voxels.

The outputs were quantified as projection signal volume in each brain region normalized by signal volume across whole brain (excluding the injection site and target area). As soma and dendrites of labeled neurons contributed a lot of signals in the injection site, we excluded the injection site for more accuracy.

### Visualization and statistical analysis

The Amira software (v6.1.1, FEI) and Imaris software (v9.5.0, Bitplane) were used to visualize the inputs and outputs of glutamatergic and GABAergic neurons in the DR and MR. To compare the inputs to glutamatergic and GABAergic neurons in the DR and MR across brain regions, we performed one-way ANOVA followed by multiple comparisons with Tukey’s test. To compare the outputs of glutamatergic and GABAergic neurons in the same nucleus across brain regions, we performed one-way ANOVA. To quantify the similarities of input and output patterns, we calculated Pearson’s correlation coefficients. To explore the similarities and variances of inputs/outputs of brain regions connected with glutamatergic and GABAergic neurons in the DR and MR, we performed correlation analysis and hierarchical cluster analysis. These processes were performed using MATLAB (v2017a, MathWorks) and Python 3.6.4. To compare the whole-brain inputs across all samples, hierarchical clustering and bootstrapping were performed using pvclust that is a package of R (Suzuki and Shimodaira, 2006). All histograms were generated using GraphPad Prism (v.6.0, GraphPad).

## Acknowledgements

We thank H.Ni, M.Ren, X.Wang for help with experiments, data analysis and constructive comments. We thank the members of MOST group of Britton Chance Center for Biomedical Photonics and HUST-Suzhou Institute for Brainsmatics for help with experiments and data acquisition. This work was financially supported by the National Natural Science Foundation of China (Grant Nos. 91749209, 61890953, 91827901), the Science Fund for Creative Research Group of China (Grant No.61721092).

## Competing interests

The authors declare no competing interests.

## Author contributions

Qingming Luo and Hui Gong conceived and designed the study. Xiangning Li performed the tracing experiments and sample preparation. Lei Deng, Tao Jiang and Jing Yuan performed the whole-brain data acquisition. Zhengchao Xu, Zhao Feng, Mengting Zhao, Xueyan Jia, Wu Chen and Anan Li performed the imaging processing and visualization. Zhengchao Xu, Qingtao Sun and Pan Luo performed data analysis and result interpretation. Zhengchao Xu, Anan Li, Hui Gong and Qingming Luo drafted the manuscript.

## Supplementary Figures

**Figure 1— figure supplement 1.**
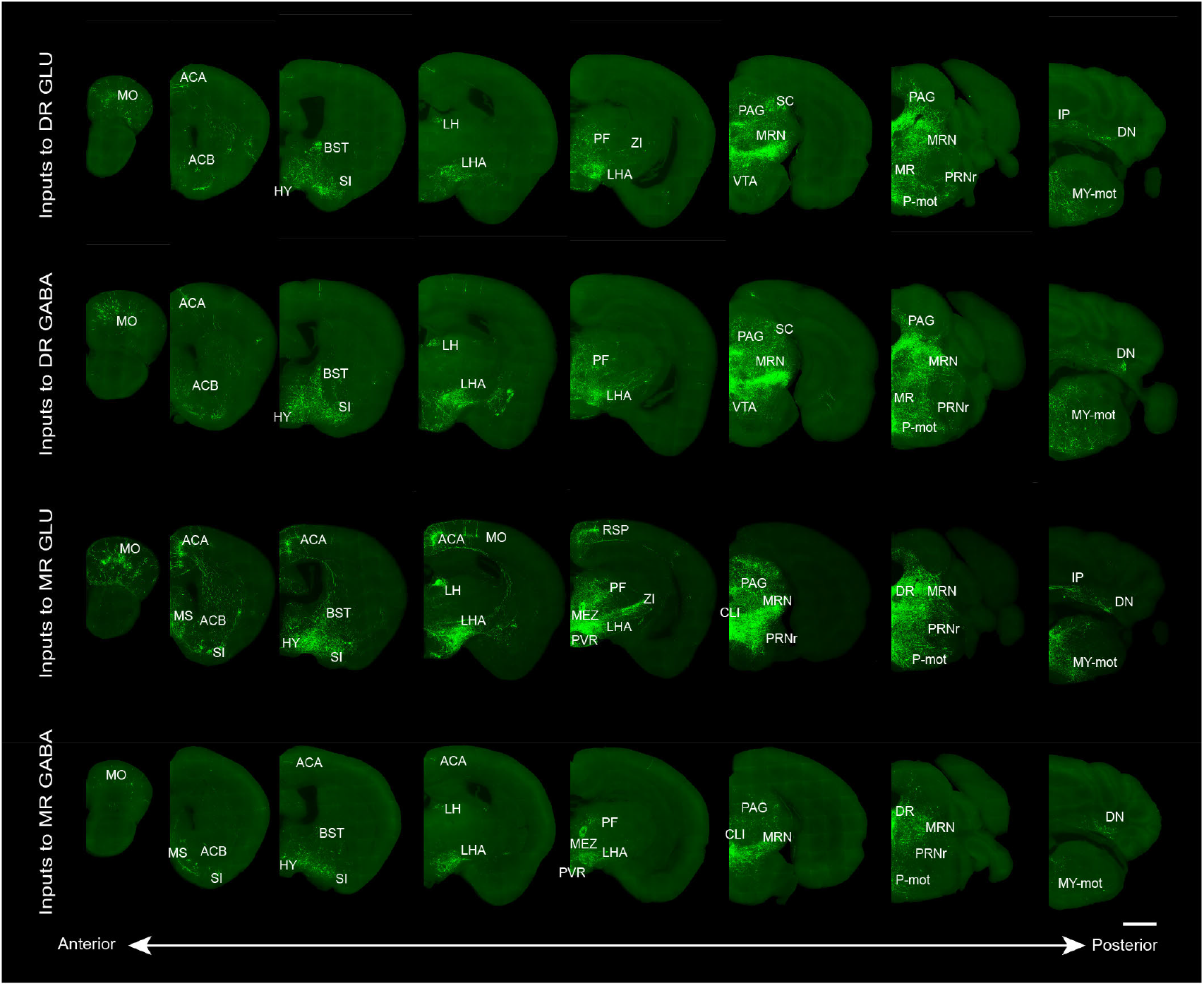
Representative images showing whole-brain inputs to the glutamatergic and GABAergic neurons in the DR and MR. The images are from maximum intensity projections of the coronal sections of representative samples. The projections were 50μm thick. Scale bar, 1 mm. The details of abbreviations for brain regions see Supplementary File 1.

**Figure 2— figure supplement 1.**
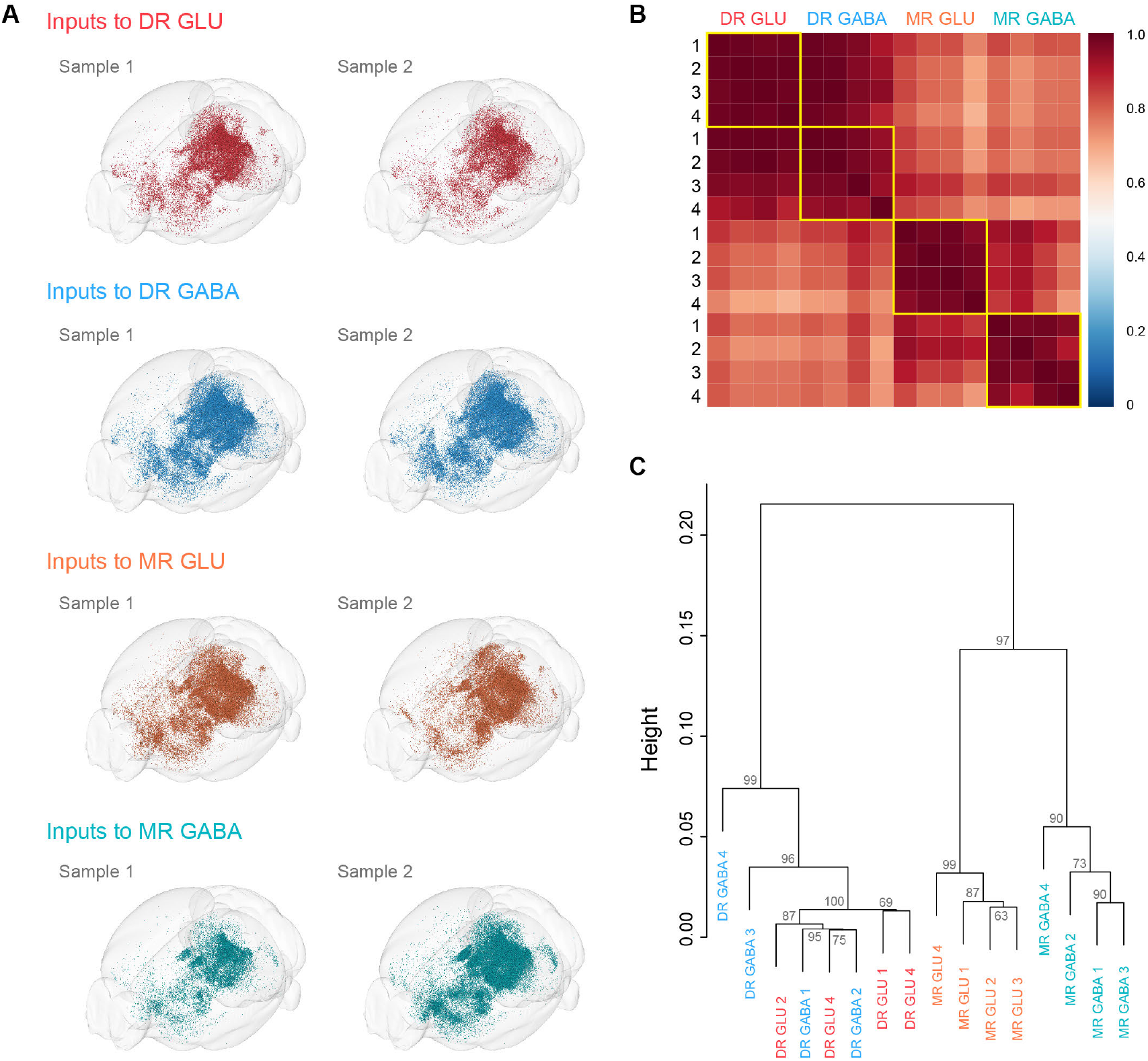
Visualization and comparison of whole-brain inputs across samples. (**A**) Visualization of whole-brain inputs to glutamatergic and GABAergic neurons in the DR and MR. The results of two individual samples of the same group of neurons demonstrate the reliability of input patterns. (**B**) Pearson’s correlation coefficients matrix of quantitative whole-brain inputs across individual samples. (**C**) Hierarchical clustering and bootstrapping based on the quantified whole-brain inputs. The approximately unbiased value shown for each branch indicates the confidence that the cluster is supported by the data.

**Figure 4— figure supplement 1.**
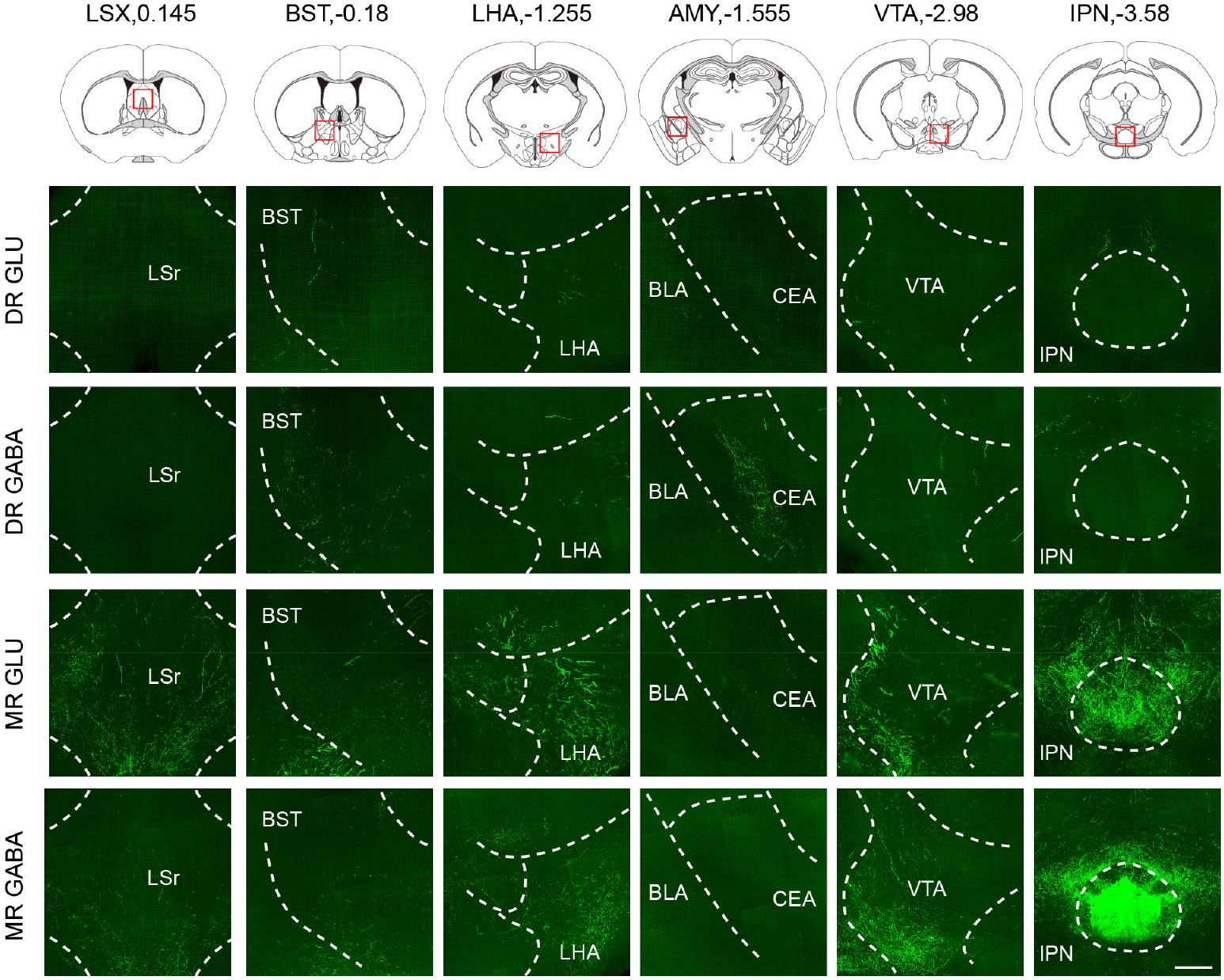
Representative images showing projections from glutamatergic and GABAergic neurons in the DR and MR. The images are from maximum intensity projections of the coronal sections of representative samples. The projections were 100 μm thick. Scale bar, 200 μm. LSX, lateral septal complex; BST, bed nuclei of the stria terminalis; LHA, lateral hypothalamic area; CEA, central amygdalar nucleus; BLA, basolateral amygdalar nucleus; VTA, ventral tegmental area; IPN, interpeduncular nucleus.

**Figure 4— figure supplement 2.**
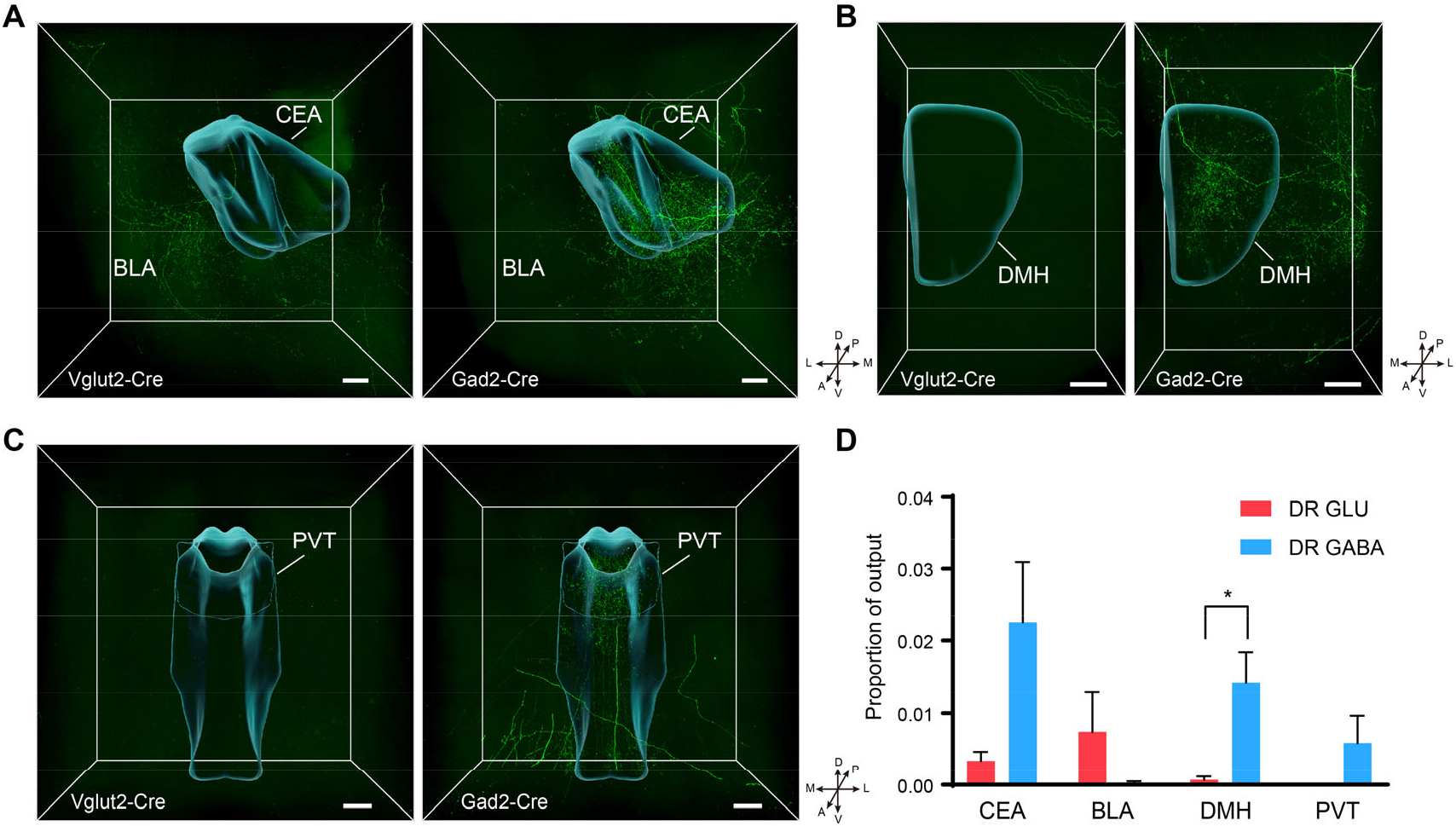
Comparison of outputs from glutamatergic and GABAergic neurons in the DR. (**A**) Three-dimensional rendering of projections in the amygdala from glutamatergic and GABAergic neurons in the DR. Scale bar, 200 μm. (**B**) Three-dimensional rendering of projections in the DMH from glutamatergic and GABAergic neurons in the DR. Scale bar, 200 μm. (**C**) Three-dimensional rendering of projections in the PVT from glutamatergic and GABAergic neurons in the DR. Scale bar, 200 μm. (**D**) Quantification and comparison of the proportion of outputs in the target area. One-way ANOVA, *p < 0.05. Data are shown as mean ± s.e.m., n=4 per group. A, anterior; P, posterior; M, medial; L, lateral; D, dorsal; V, ventral. CEA, central amygdalar nucleus; BLA, basolateral amygdalar nucleus; DMH, dorsomedial nucleus of the hypothalamus; PVT, Paraventricular nucleus of the thalamus;

**Figure 5— figure supplement 1.**
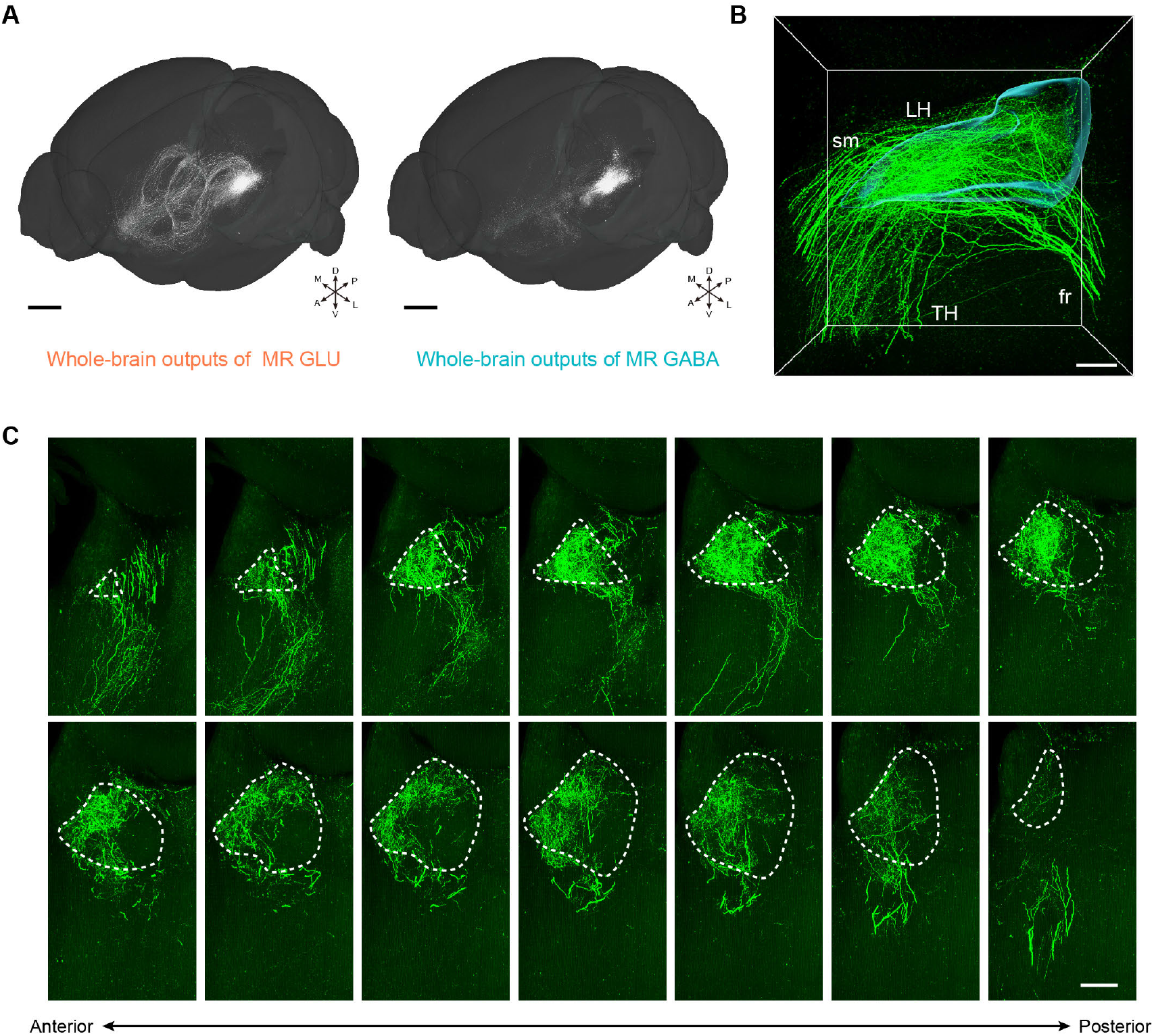
Representative outputs of glutamatergic and GABAergic neurons in the MR. (**A**) Whole-brain outputs of glutamatergic and GABAergic neurons in the MR of representative samples. Scale bar, 1 mm. (**B**) Enlarged view of three-dimensional rendering of projections in the LH (left hemisphere) from MR glutamatergic neurons. Scale bar, 200 μm. (**C**) Images showing the projections in the LH (left hemisphere) from MR glutamatergic neurons. The images are from maximum intensity projections of the coronal sections of a representative sample. The projections were 100 μm thick. Scale bar, 200 μm. A, anterior; P, posterior; M, medial; L, lateral; D, dorsal; V, ventral.

**Figure 6— figure supplement 1.**
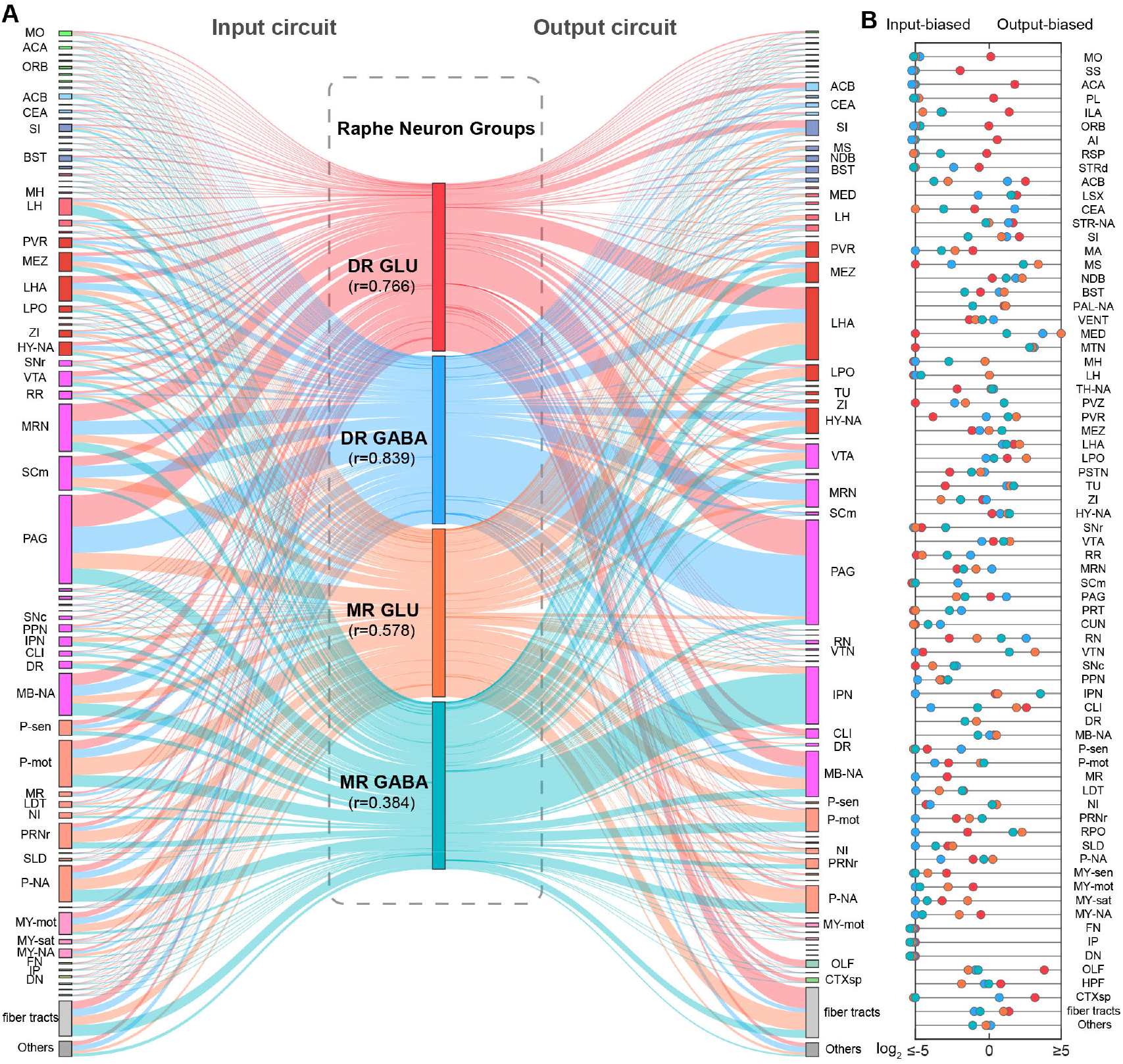
Connectivity pattern of whole-brain inputs and outputs of glutamatergic and GABAergic neurons in DR and MR. (**A**) Schematic and similarity of whole-brain inputs and outputs of glutamatergic and GABAergic neurons in the DR and MR. The line width reflects proportion. The colors reflect the neurons types. DR glutamatergic and GABAergic neurons and their connections are respectively reflected by red and blue. MR glutamatergic and GABAergic neurons and their connections are respectively reflected by orange and turquoise. r: Pearson’s correlation coefficients reflecting the similarity of whole-brain inputs and outputs of the same group of neurons. (**B**) Reciprocity between whole-brain inputs and outputs of glutamatergic and GABAergic neurons in the DR and MR. The scatter represents the ratio of output and input across brain regions, where colors reflect the neuron types as in (**A**). The details of abbreviations for brain regions see Supplementary File 1.

**Supplementary File**

**Supplementary File 1. Nomenclature and abbreviations of brain regions**

**Supplementary File 2. Quantification and comparison of whole-brain inputs to glutamatergic and GABAergic neurons in the DR and MR**

**Supplementary File 3. Quantification and comparison of whole-brain outputs of glutamatergic and GABAergic neurons in the DR and MR**

